# An RNAi screen for conserved kinases that enhance microRNA activity after dauer in *C. elegans*

**DOI:** 10.1101/2023.10.17.562753

**Authors:** Himal Roka Pun, Xantha Karp

**Affiliations:** Department of Biology, Central Michigan University, Mount Pleasant, MI 48859; Biochemistry, Cell and Molecular Biology Program Central Michigan University, Mount Pleasant, MI 48859

**Keywords:** Diapause, dauer, *C. elegans*, microRNA, Argonaute, kinase

## Abstract

Gene regulation in changing environments is critical for maintaining homeostasis. Some animals undergo a stress-resistant diapause stage to withstand harsh environmental conditions encountered during development. MicroRNAs (miRNAs) are one mechanism for regulating gene expression during and after diapause. MicroRNAs downregulate target genes post-transcriptionally through the activity of the miRNA-Induced Silencing Complex (miRISC). Argonaute is the core miRISC protein that binds to both the miRNA and to other miRISC proteins. The two major miRNA Argonautes in the *C. elegans* soma are ALG-1 and ALG-2, which function partially redundantly. Loss of *alg-1 (alg-1(0))* causes penetrant developmental phenotypes including vulval defects and the reiteration of larval cell programs in hypodermal cells. However, these phenotypes are essentially absent if *alg-1(0)* animals undergo a diapause stage called dauer. Levels of the relevant miRNAs are not higher during or after dauer, suggesting that activity of the miRISC may be enhanced in this context. To identify genes that are required for *alg-1(0)* mutants to develop without vulval defects after dauer, we performed an RNAi screen of genes encoding conserved kinases. We focused on kinases because of their known role in modulating miRISC activity. We found RNAi knockdown of four kinase-encoding genes, *air-2*, *bub-1*, *chk-1,* and *nekl-3*, caused vulval defects and reiterative phenotypes in *alg-1(0)* mutants after dauer, and that these defects were more penetrant in an *alg-1(0)* background than in wild type. Our results implicate these kinases as potential regulators of miRISC activity during post-dauer development in *C. elegans*.

## Introduction

The regulation of gene expression in response to changing environments is critical to maintaining homeostasis. One mechanism for modulating gene expression in different contexts is microRNA-mediated gene silencing. MicroRNAs (miRNAs) are ∼22 nucleotide non-coding RNAs that regulate mRNA transcripts post-transcriptionally via the <u>miR</u>NA Induced <u>S</u>ilencing <u>C</u>omplex (miRISC). MiRNAs bind to the 3’UTR of the target mRNA via imperfect complementarity, thereby recruiting miRISC. Argonaute is the core miRNA-interacting protein that in turn binds with other protein cofactors (Bartel 2018). The protein factors in miRISC inhibit translation and cause destabilization of the transcript, thus silencing the expression of the target (Jonas and Izaurralde 2015; Bartel 2018).

Changes in miRNA activity can influence gene expression in a variety of physiological or environmental contexts (Leung and Sharp 2010; Galagali and Kim 2020). When confronted with adverse environments, some animals can enter a stress-resistant and developmentally arrested diapause stage (Hand *et al*. 2016). MicroRNAs have been implicated in the regulation of diapause in diverse animal species, including insects, killifish, and nematodes (Alvarez-Saavedra and Horvitz 2010; Karp *et al*. 2011; Meuti *et al*. 2018, Reynolds 2019). Although changes in miRNA levels have been documented before, during, and after diapause, little is known about changes in miRISC activity and how they may impact development in the diapause life history.

*C. elegans* larval development is a useful model system to explore the modulation of miRISC activity during diapause. *C. elegans* larvae develop through four larval stages (L1-L4) before becoming adults (Byerly *et al*. 1976). Larval development occurs continuously when the environment is favorable for reproduction. In unfavorable environments, larvae can pause their development after the second larval molt in the stress-resistant dauer diapause stage (Cassada and Russell 1975). If dauer larvae encounter favorable environmental conditions, they can recover and resume development through post-dauer larval stages that are developmentally equivalent to those occurring in continuously developing larvae (Liu and Ambros 1991; Euling and Ambros 1996).

One developmental system regulated by miRNAs is the hypodermal stem-cell-like seam cells. Seam cells divide in a particular pattern and sequence at each larval stage. At adulthood, seam cells cease dividing and differentiate (Sulston and Horvitz 1977; Ambros and Horvitz 1984). Larval vs. adult seam programs are regulated by heterochronic genes that encode miRNAs and their targets (Ambros and Horvitz 1984; Rougvie and Moss 2013; Galagali and Kim 2020). The loss of heterochronic miRNAs or impaired function of miRISC proteins causes a reiterative phenotype whereby the pattern and sequence of seam cell divisions appropriate to earlier developmental stages is reiterated in later stages (Ambros and Horvitz 1984; Reinhart *et al*. 2000; Grishok *et al*. 2001; Abbott *et al*. 2005).

ALG-1 and ALG-2 are the best characterized somatic miRNA-specific Argonaute proteins in *C. elegans* and act partially redundantly (Grishok *et al*. 2001; Tops *et al*. 2006; Vasquez-Rifo *et al*. 2012). In addition, other Argonautes including ALG-5 and RDE-1 can also bind miRNAs (Seroussi *et al*. 2023). While loss of both *alg-1* and *alg-2* results in embryonic lethality, loss of *alg-1* alone *(alg-1(0))* causes a reiterative phenotype, suggesting ALG-2 and the other Argonautes are not sufficient to mediate the activity of the heterochronic miRNAs (Grishok *et al*. 2001; Zinovyeva *et al*. 2014). Strikingly, this phenotype is strongly suppressed in post-dauer *alg-1(0)* mutant animals (Karp and Ambros 2012). This finding suggests that miRNA activity is enhanced after dauer, such that *alg-2* activity becomes sufficient. Prior work has demonstrated that levels of heterochronic miRNAs are the same or reduced in post-dauer larvae compared to continuously developing larvae, suggesting that the enhanced miRNA activity may arise from modulation of miRISC (Karp *et al*. 2011; Karp and Ambros 2012).

Kinases are good candidates to mediate miRISC enhancement after dauer because they are known modulators of miRISC activity. In *C. elegans* and other animals, kinases phosphorylate Argonaute and other miRISC components, thereby altering miRISC activity (Wilczynska and Bushell 2015; Frédérick and Simard 2022). To identify conserved kinases that may enhance miRNA activity after dauer, we carried out an RNAi screen for conserved kinase-encoding genes that are necessary for the post-dauer suppression of *alg-1(0)* reiterative phenotypes. We found that *air-2, bub-1, chk-1,* and *nekl-3* are required to prevent vulval defects and reiterative phenotypes in *alg-1(0)* mutants after dauer. Furthermore, RNAi of these genes causes more penetrant phenotypes in an *alg-1* background than in wild-type, suggesting that the kinases they encode may enhance miRNA function after dauer.

## Methods

### *C. elegans* strain and maintenance

All strains were maintained on Nematode Growth Medium (NGM) plates seeded with *E. coli* strain OP50 at 15°C or 20°C (Brenner 1974). Strains used in the study were: XV88 *daf-7(e1372); maIs105[col-19p::gfp]; alg-1(gk214),* VT1777 *daf-7(e1372); maIs105,* XV85 *alg-2(ok302); daf-7(e1372); maIs105,* and VT1274 *alg-2(ok302); maIs105*.

### RNAi

A list of genes encoding kinases conserved between *C. elegans* and humans was obtained from (Deng *et al*. 2019). Their list was generated using the Ortholist tool (Shaye and Greenwald 2011; Kim *et al*. 2018). A complete list of RNAi clones used, and their sources is provided in Table S1. RNAi bacteria were grown in LB media containing 50µL/mL carbenicillin shaking overnight at 37°C. RNAi bacteria were then seeded on NGM plates containing 200µg/mL IPTG and 50µg/mL carbenicillin (“RNAi plates”). For the primary screen, 12-well plates were used, and for all other experiments, 60mm plates were used. The primary screen was conducted in duplicate. RNAi clones that caused reiterative alae defects in post-dauer adults were verified by Sanger sequencing using the M13F primer (Eurofins Genomics).

#### Embryo Isolation

Gravid adult hermaphrodites were treated with a bleach solution (0.4% sodium hypochlorite dissolved in 1M NaOH) for two 2-minute incubations at room temperature. The embryos were washed with sterile water and then added to RNAi plates.

#### Dauer Induction

The *daf-7(e1372)* allele was used to induce dauer formation (Vowels and Thomas 1992; Karp 2018). Embryos were incubated on RNAi plates at 24°C for 48 – 50 hours, corresponding to the time just after the molt into dauer (Karp 2018).

#### Dauer recovery

For the primary screen, dauer recovery was induced by washing dauer larvae off the RNAi plates with sterile water, adding the dauer larvae to fresh RNAi plates, and shifting the worms to 20°C. For all other experiments, dauer larvae were selected by incubating worms with 1% (w/v) SDS (sodium dodecyl sulfate) for 30 minutes at room temperature (Cassada and Russell 1975; Karp 2018). The dauer larvae were then washed with sterile water, added to fresh RNAi plates, and shifted to 20°C to induce dauer recovery. Post-dauer larvae were incubated at 20°C for 48 hours to obtain young post-dauer adults.

#### New RNAi clones

RNAi clones for *air-2, chk-1,* and *bub-1* were created based on the protocol described in (Kamath and Ahringer 2003). Briefly, primers were designed to amplify regions of these genes that were distinct from those contained in the Ahringer RNAi clones. The amplified sequences were cloned into the L4440 vector by TA cloning. The resulting plasmids were sequenced, then transformed into HT115 bacteria. Primers used for cloning are listed below.

*air-2* forward: 5’-TACTCCACAGAAGGGAGGGT-3’

*air-2* reverse: 5’-ATGTTGGCCACTAAGCTGAAATC-3’

*chk-1* forward: 5’-GGCGGAGAGACAGAATGCTT-3’

*chk-1* reverse: 5’-CCGAGTGCTCCACATTGACT-3’

*bub-1* forward: 5’ CCGTCGACATGTGGTCTTGA-3’

*bub-1* reverse: 5’-GAGGTTTGCGTCACTGGAGA-3’

#### Microscopy

For the primary screen, bursting and/or protrusion of the vulva (Rup or Pvl phenotypes) of the post-dauer adults were assessed using a dissecting microscope (Zeiss Stereo V12 fitted with M2 Bio for fluorescence). In the secondary screen, the *alg-1(0)* mutant phenotypes were assessed using a Zeiss AxioImager D2 compound microscope. Post-dauer animals were immobilized using 0.1M levamisole on 2% agarose pads. DIC and fluorescence images were taken using an AxioCam MRm Rev 3 camera and ZEN 3.2 software. GFP was visualized with a high-efficiency GFP shift-free filter at 63x with an exposure time of 9ms.

## Results and discussion

### RNAi screen of the conserved kinome to identify miRISC regulators acting after dauer

If *C. elegans* larvae lacking *alg-1* develop continuously from embryo to adult, the adults display reiterative phenotypes (Grishok *et al*. 2001; Zinovyeva *et al*. 2014). In contrast, these phenotypes are suppressed if *alg-1(0)* mutant larvae develop through the dauer diapause stage (Karp and Ambros 2012). We hypothesized that modulation of miRISC activity in post-dauer animals could account for this difference. Since kinases are known to modulate miRISC in various physiological and environmental contexts (Wu *et al*. 2011; Alessi *et al*. 2015; Olejniczak *et al*. 2018), we designed an RNAi screen of all of the conserved kinases in *C. elegans* (Kim *et al*. 2018; Deng *et al*. 2019). If post-dauer *alg-1(0)* mutants display reiterative phenotypes after depletion of a particular kinase, that kinase is a candidate for modulating miRISC function in post-dauer animals.

To perform our RNAi screen, we used an *alg-1(0)* strain that contained the temperature-sensitive *daf-7(e1372)* allele to induce dauer formation at 24°C and allow recovery at lower temperatures (Vowels and Thomas 1992; Karp 2018). A *col-19p::gfp* transgene was also included to allow analysis of that aspect of adult cell fate (Liu et al. 1995). Moving forward, unless otherwise specified, it should be presumed that all strains described here contain *daf-7(e1372); maIs105[col-19p::gfp]* in the background. Embryos were added to RNAi plates and incubated at 24°C for 2 days, or until just after dauer formation. Dauer larvae were washed off with water, transferred to fresh RNAi plates, and shifted to 20°C to promote recovery. Post-dauer *alg-1(0)* adults were then screened for the vulval defects associated with compromised miRISC function (Figure 1A). We used *lacZ* RNAi as a negative control and *kin-3* RNAi as a positive control because we previously showed that RNAi of *kin-3* caused reiterative phenotypes in post-dauer *alg-1(0)* mutants (Alessi *et al*. 2015). *kin-3* encodes the catalytic subunit of Casein Kinase II (CK2), and CK2 phosphorylates the miRISC proteins ALG-1 and CGH-1 to promote miRISC function (Alessi *et al*. 2015; Shah *et al*. 2023). We used a previously published list of the 247 *C. elegans* kinases with human orthologs (Kim *et al*. 2018; Deng *et al*. 2019). Some kinases are represented by more than one RNAi clone, giving us 286 RNAi clones in total (Table S1).

**Fig. 1.**
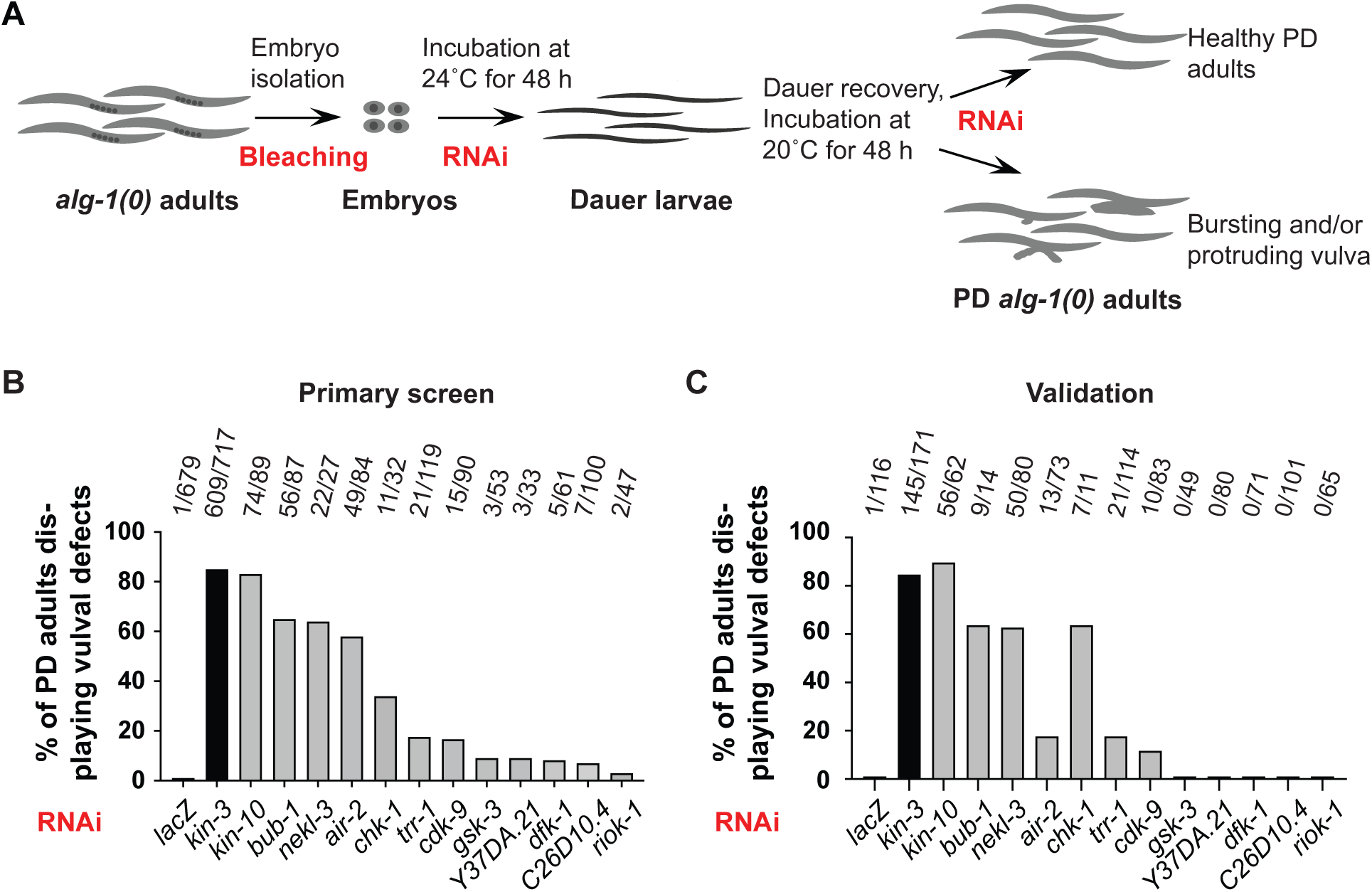
Kinases identified in a primary screen for enhancement of an *alg-1(0)* phenotype after dauer. **A.** Strategy for RNAi screening. The full genotype of the strain used for screening was *daf-7(e1372); maIs105[col-19p::gfp]; alg-1(gk214).* The *daf-7(e1372)* allele was used to induce dauer formation at 24°C and allow recovery at lower temperatures. See Methods for details. **B.** RNAi clones that produced vulval defects (Rup and/or Pvl phenotypes) in the primary screen. The screen was performed in duplicate and clones that produced vulval defects in both plates are shown here. Table S1 shows data for all clones that produced vulval defects in at least one well. The remaining clones produced no vulval defects in either well (≥20 adults scored per well). Numbers in B and C indicate the number of post-dauer (PD) adults displaying vulval defects over the total PD adults scored. **C.** The 12 hits from the primary screen were retested for vulval defects. Seven genes caused bursting and/or protrusion of the vulva in post-dauer *alg-1(0)* adults. Among these 7 genes, 6 were novel *alg-1* interactors.

In our primary screen, performed in duplicate, we used a dissecting microscope to look for vulval defects in *alg-1(0)* post-dauer adults. Vulval defects included both bursting (Rup) and protruding vulva (Pvl) phenotypes. We focused on vulval defects because these phenotypes are easily visible in the dissecting microscope. As expected, we saw almost no vulval defects after treatment with *lacZ* RNAi (Figure 1B). By contrast, *kin-3* RNAi caused a high penetrance of vulval defects (Figure 1B). We found 12 additional conserved kinases that caused similar phenotypes. We observed a broad range of penetrance of vulval defects which could be due to different degrees of requirement for vulval development and/or variable effectiveness of RNAi.

To ensure our results were reproducible, we re-tested the 12 hits from the primary screen. In addition, in this validation experiment, we selected dauer larvae by treatment with 1% SDS (Cassada and Russell 1975). Treatment with SDS ensured that the adults we screened were indeed post-dauer and not displaying vulval defects because the larvae had escaped dauer formation. Upon retesting, seven genes displayed vulval defects (Figure 1C). One of these genes, *kin-10*, encodes the regulatory subunit of the of CK2 protein and was therefore expected (Alessi *et al*. 2015; Shah *et al*. 2023). We therefore focused on the remaining six kinase-encoding genes.

### Five kinase-encoding genes contribute to adult alae formation in *alg-1(0)* post-dauer adults

In addition to vulval defects, *alg-1(0)* mutant adults display gapped or missing alae after continuous development (Grishok *et al*. 2001; Vasquez-Rifo *et al*. 2012). This reiterative phenotype occurs in heterochronic mutants because the underlying seam cells repeat larval cell programs (Ambros and Horvitz 1984). In addition to gapped or missing alae, *alg-1(0)* animals that develop continuously can display other defects in alae formation, including small breaks and growth that deviates from the straight line observed in wild-type adults (Alessi *et al*. 2015; Shah *et al*. 2023). Furthermore, the loss of *alg-1* can reduce the expression of the adult cell fate marker *col-19p::gfp,* albeit at low penetrance (Zinovyeva *et al*. 2015). If the kinases we identified were important for miRNA function we would expect depletion of these genes to cause alae defects and possibly reduced *col-19p::gfp* in the hypodermis of post-dauer *alg-1(0)* adults.

To test the hypothesis that these kinases are required to prevent reiterative phenotypes in post-dauer *alg-1(0)* adults, we performed RNAi on the six hits and looked at adult alae formation and *col-19p::gfp* expression with DIC and fluorescence microscopy. We found that knocking down five of the six genes with RNAi resulted in penetrant alae defects (Figure 2). The most penetrant alae phenotypes were caused by *bub-1* RNAi. RNAi of *cdk-9* produced no reiterative phenotypes and occasional other alae defects. While *lacZ* RNAi produced no alae defects at all, *cdk-9* RNAi was not significantly different from the *lacZ* control. In contrast to the alae defects produced by RNAi, *col-19p::gfp* expression was not reduced by RNAi of any of the six candidate genes and only moderately downregulated in our *kin-3* positive control (Figure S1). Uncoupling of alae defects and *col-19p::gfp* expression has been previously observed in some mutant backgrounds (Hada *et al*. 2010; Hansen *et al*. 2022).

**Fig. 2.**
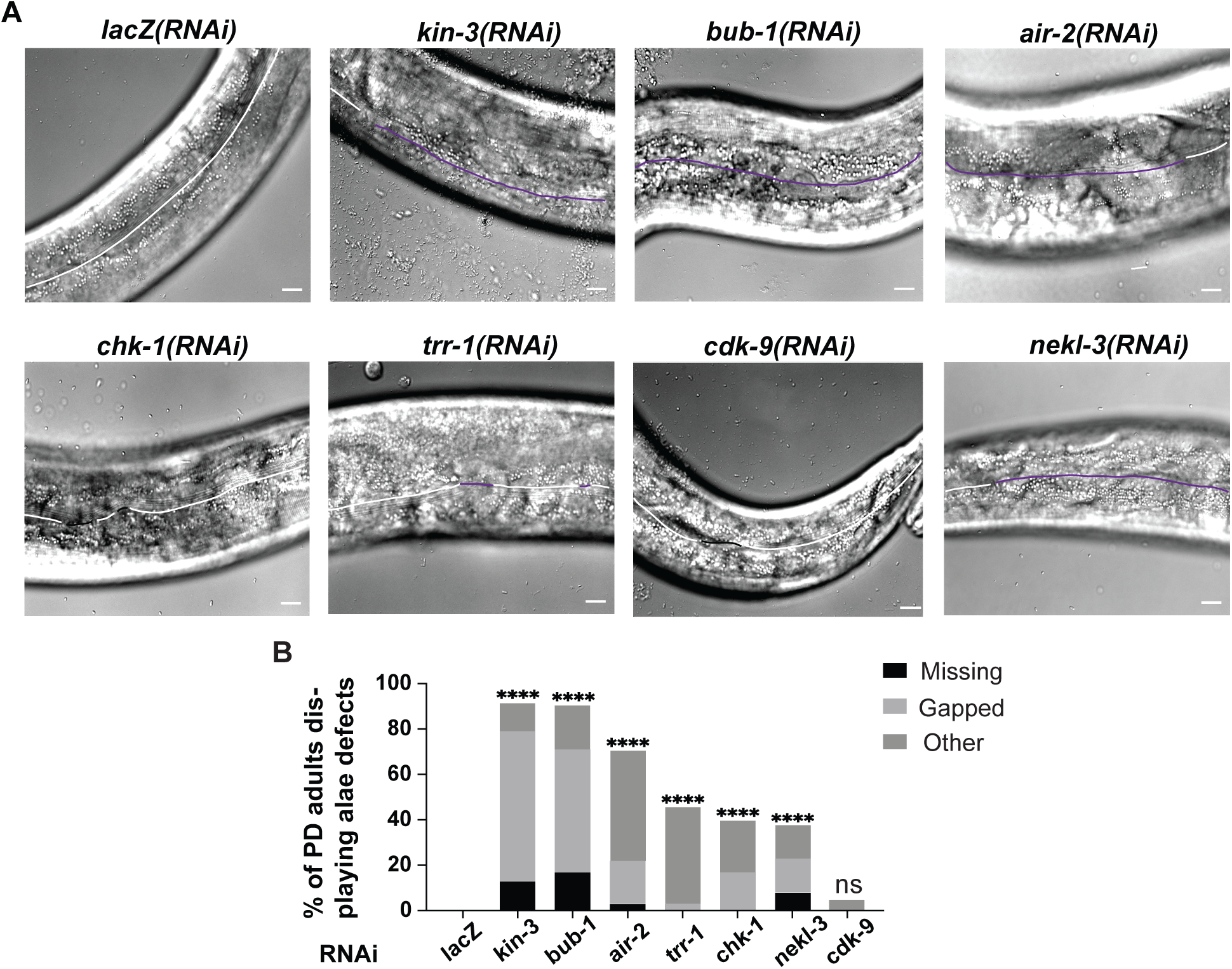
RNAi of five kinase-encoding genes produces defects in adult alae formation in post-dauer *alg-1(0)* mutants. **A**. Comparison of alae phenotypes in post-dauer *daf-7(e1372); col-19p::gfp; alg-1(0)* adults. A compound microscope with DIC optics was used to visualize the presence of alae in post-dauer adult animals. White lines represent complete alae, purple lines represent gapped alae, and black lines show other alae defects including small breaks or curved or swirly alae. **B**. Quantification of alae defects. **** = *P* <0.0002; ns = *P* > 0.05, Fisher Exact Test.

### RNAi of some kinases produces a stronger phenotype in a miRISC-sensitized background and during post-dauer development

Vulval defects can be caused by the depletion of genes that regulate processes other than miRISC activity. For example, loss of genes involved in regulation of uterine development can produce similar phenotypes (Seydoux and Greenwald 1989). If the kinases identified in our screen enhance miRISC activity after dauer, we would expect that RNAi of these kinases would produce a stronger phenotype in *alg-1(0)* adults than when *alg-1* was wild type. To test this hypothesis, we performed RNAi on *alg-1(0)* and *alg-1(+)* animals in parallel. RNAi of *bub-1, air-2, nekl-3,* and *chk-1* each displayed a more penetrant phenotype in the miRISC-compromised background (Figure 3A). RNAi of *trr-1* or *cdk-9* caused a low-penetrance phenotype in both backgrounds, and there was no significant difference between the backgrounds (Figure 3A). While these data do not rule out the involvement of any of these genes in regulating miRISC activity, they provide higher confidence in *bub-1, air-2, nekl-3,* and *chk-1* than *trr-1* or *cdk-9*.

**Fig. 3.**
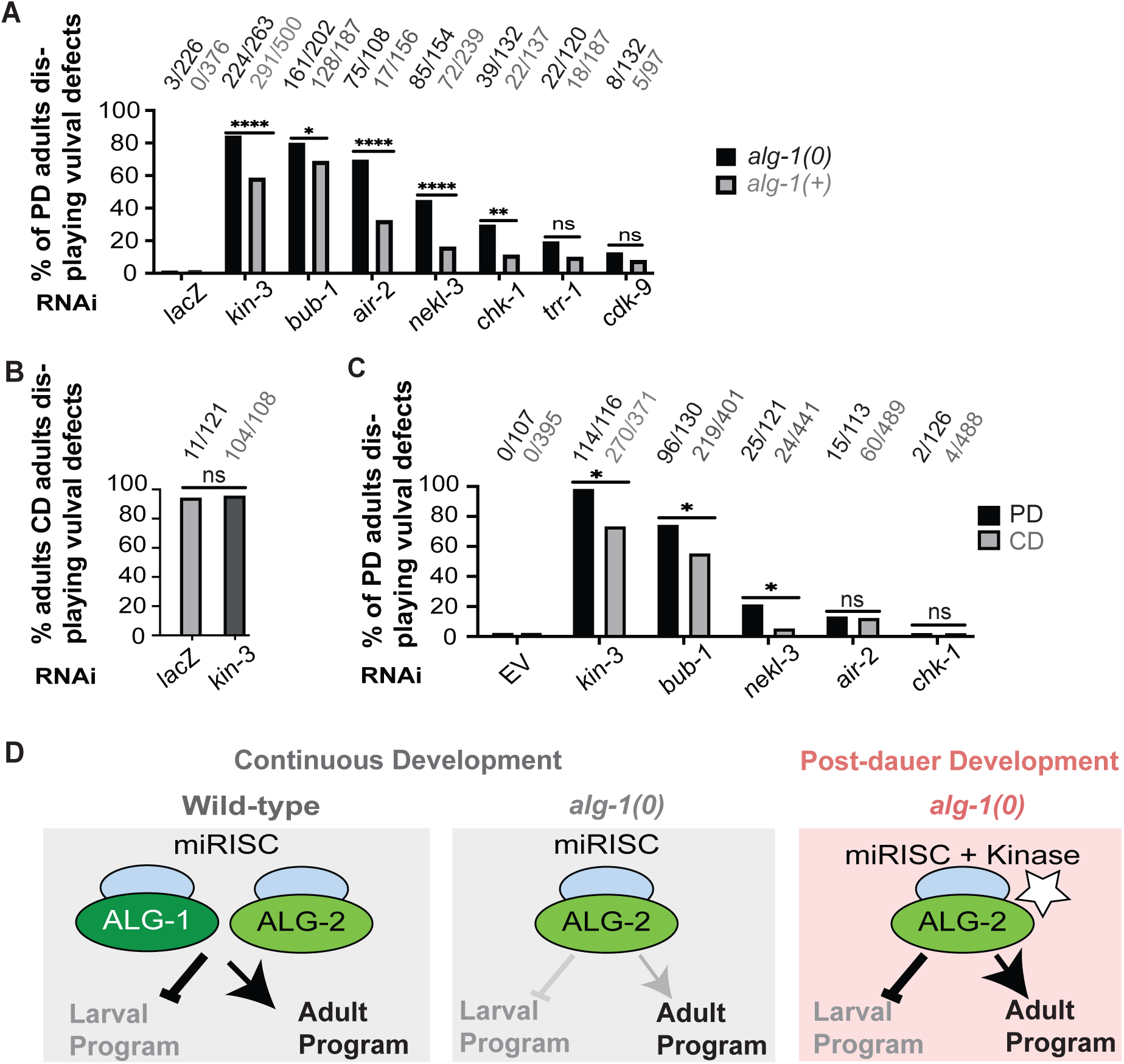
RNAi of some kinases caused more penetrant phenotypes after dauer and when miRISC was compromised. **A.** Vulval defects in post-dauer adults with wild-type miRISC were compared to animals with compromised miRISC function *(alg-1(0)* mutants). Both strains contained *daf-7(e1372); col-19p::gfp* in the background. **B.** During continuous development, the penetrance of vulval defects in *alg-1(0)* adults is high such that no further enhancement is seen when *kin-3* is knocked down. **C.** Vulval defects in *alg-2(0)* young adults after dauer (PD = post-dauer) or after continuous development (CD). In B and C, dauer was induced by *daf-7(e1372),* whereas continuously developing animals were wild-type for *daf-7*. *p-*value >0.05 (ns), <0.0332 (*), <0.0021 (**), <0.0002 (****), Fisher exact test. Empty vector (EV) was used as a negative control. **D)** Model for role of miRISC activity in different developmental contexts to control larval vs. adult seam cell developmental programs. Note that other Argonautes could act in addition to or in place of ALG-2.

Since *alg-1(0)* mutants display vulval defects during continuous development but not after dauer, we next wondered whether the kinases we identified play a greater role in preventing vulval defects in post-dauer hermaphrodites than in those that develop continuously, particularly when miRISC is compromised. We were unable to test this question using *alg-1(0)* mutants because they display penetrant phenotypes during continuous development (Figure 3B). However, animals lacking *alg-2* appear superficially wild-type in both life histories (Grishok *et al*. 2001; Karp and Ambros 2012). We therefore compared phenotypes in each life history following RNAi of *bub-1, nekl-3, air-2*, and *chk-1* in an *alg-2(0)* mutant background. We found that RNAi of *bub-1* and *nekl-3* both produced statistically more penetrant phenotypes after dauer (Figure 3C), consistent with these kinases playing a larger role in post-dauer development. By contrast, RNAi of *air-2* and *chk-1* produced low-penetrance phenotypes in both life histories in the *alg-2(0)* background (Figure 3C). While in this latter case there was no statistical difference between post-dauer and continuously developing animals, observing a decrease in penetrance in adults that had developed continuously would be difficult given the low penetrance phenotype in post-dauer adults. Thus, we are unable to draw strong conclusions about the relative role of *air-2* and *chk-1* in post-dauer vs. continuous development.

Finally, we performed RNAi of *bub-1, air-2, nekl-3,* and *chk-1* using independent clones that target different regions of each gene than the region targeted in the original clones. We observed vulval defects using each set of clones, albeit at reduced penetrance (Figure S2). The percent of vulval defects observed for each clone was statistically different from the empty vector control, except the newly created *chk-1* clone (P = 0.088). Both original and newly created *chk-1* clones produced low-penetrance phenotypes, whereas the negative control produced no vulval defects out of 226 adults examined.

### Kinases identified in this screen

We have identified four kinases that, when depleted, reproducibly enhanced two different *alg-1(0)* phenotypes in post-dauer animals and whose depletion caused more severe defects in a miRISC-compromised background than in wild type (Table 1). The known roles for these kinases are described below.

**Table 1:**
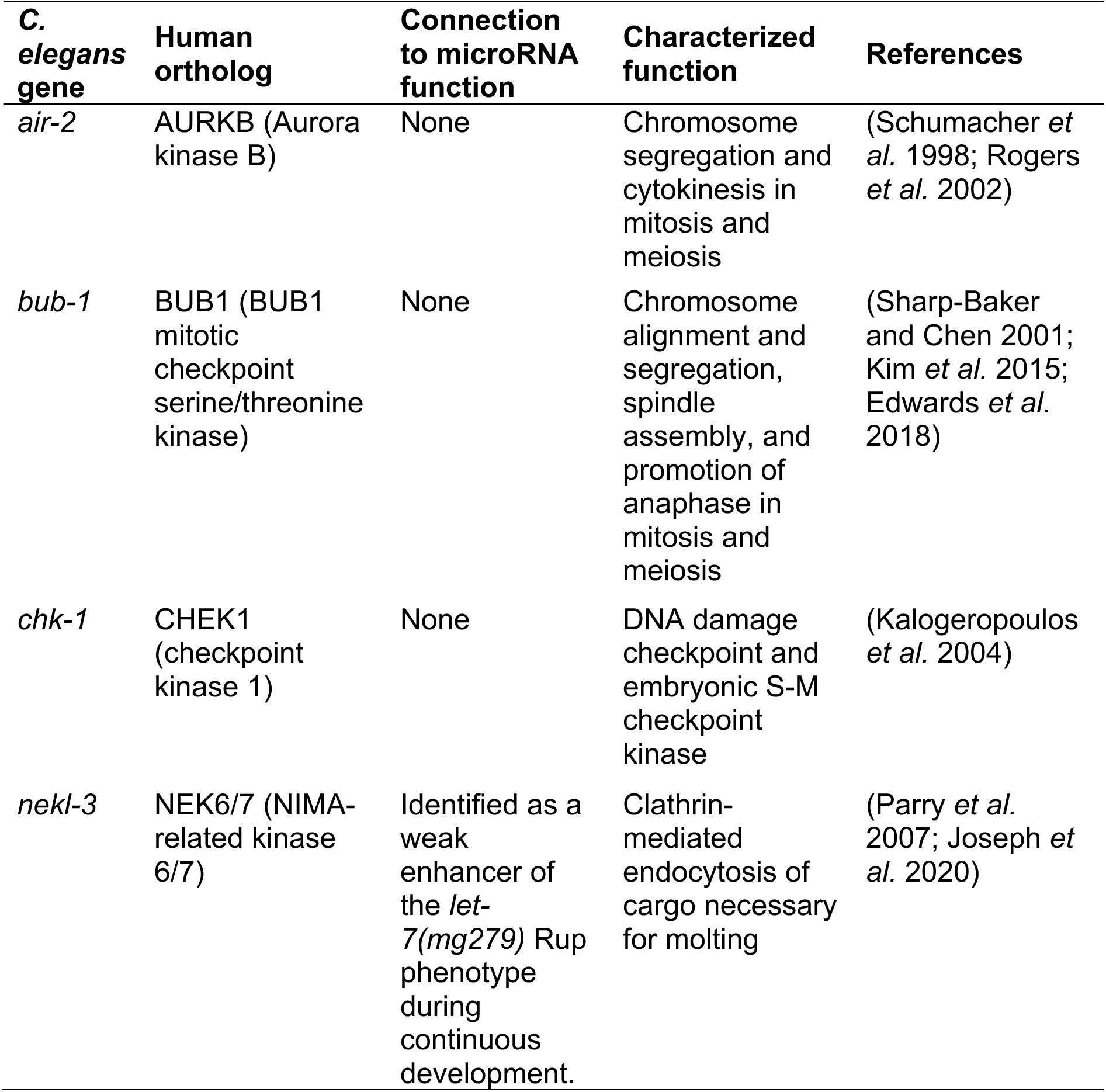
Conservation and function of genes isolated in the screen.

Among the four genes identified as the highest confidence candidates in our screen, *nekl-3* is the only one to have been previously implicated in the regulation of miRNA activity. *nekl-3* was isolated in a large-scale RNAi screen for genes that enhance the Rup phenotype of mutants with reduced activity of the *let-7* miRNA (Parry *et al*. 2007). Although *nekl-3* was a relatively weak enhancer and the screen was performed during continuous (non-dauer) development, this observation is consistent with the hypothesis that *nekl-3* can promote miRISC function. This role for *nekl-3* remains unexplored. In contrast, a role for *nekl-3* in the regulation of molting has been well characterized. *nekl-3* is required for the internalization via clathrin-mediated endocytosis of cargo important for molting (Yochem *et al*. 2015; Joseph *et al*. 2020). Specifically*, nekl-*3 is required for the L1-L2 and L2-L3 molts but not later molts during continuous development. In the dauer life history, *nekl-*3 has a reduced requirement for the L2d-dauer molt and is dispensable for post-dauer molting (Binti *et al*. 2022).

In mammals, the *nekl-3* orthologs, NEK family proteins, are best characterized for their role in mitosis and chromosome segregation (Fry *et al*. 2012). Notably, *air-2* and *bub-1* both encode conserved proteins that are critical for chromosome segregation and mitosis and meiosis across species (Schumacher *et al*. 1998; Sharp-Baker and Chen 2001; Rogers *et al*. 2002; Kim *et al*. 2015). Neither of these genes have been previously implicated in miRNA activity. Previous large-scale screens conducted during continuous development found that RNAi knockdown of *bub-1* produced Pvl and bursting phenotypes (Fraser *et al*. 2000). Loss-of-function *bub-1* mutants display a vulvaless (Vul) phenotype and a low-penetrance Pvl phenotype (Wang *et al*. 2009). However, whether these phenotypes are related to the characterized role in chromosome segregation or whether they may indicate a role for *bub-1* in regulating miRNA activity is unknown. Finally, *chk-1* encodes a serine-threonine kinase that plays a role in the checkpoint response to DNA damage and in the S-M checkpoint in embryos (Kalogeropoulos *et al*. 2004).

If the kinases we identified do regulate miRISC, this regulation could be direct or indirect. Consistent with direct regulation, *bub-1* and *nekl-3* are expressed in tissues displaying *alg-1(0)* phenotypes, including the vulva and the hypodermis (Yochem *et al*. 2015, Tarailo-Graovac *et al*. 2010). Post-embryonic expression data outside of the germline has not been described for *air-2* or *chk-1.* Furthermore, no expression data for any of the four kinases has been reported in post-dauer animals. It will be interesting to determine whether these kinases are expressed in relevant tissues in that context. In addition, core miRISC proteins including ALG-1, ALG-2, AIN-1, and AIN-2, have many phospho-sites identified by phosphoproteomic studies (Huang *et al*. 2020, Li *et al*. 2021). It is unclear whether these sites may be phosphorylated by the kinases identified in this study. There are no defined consensus sequences phosphorylated by BUB-1 and NEKL-3. There are loose consensus sequences defined for the human orthologs of AIR-2 ([RK]X[TS]) and CHK-1 (RXX[TS]) that may also apply to *C. elegans* (Sanchez *et al*. 1997; Cheeseman *et al*. 2002; Chen *et al*. 2003; Bishop *et al*. 2005; Ferrandiz *et al*. 2018). ALG-1, ALG-2, AIN-1, and AIN-2 all contain sequences that match these loose consensus sites; however, these sequences were not found to be phosphorylated in the aforementioned phosphoproteomic studies (Huang *et al*. 2020, Li *et al*. 2021). Notably, these studies were not performed in post-dauer animals. It will be interesting to explore the possibility of direct regulation of miRISC components by these kinases in future work.

## Concluding Remarks

The goal of this study was to identify kinase-encoding genes that may enhance miRISC function in post-dauer development. This study initially found six candidate genes required to prevent developmental phenotypes in post-dauer *alg-1(0)* animals. Four of these genes, *bub-1, air-2, nekl-3,* and *chk-1*, showed higher penetrance phenotypes in a miRISC-compromised background, making these four genes the most likely to affect miRISC activity after dauer. *bub-1* and *nekl-3* produced more penetrant phenotypes after dauer than during continuous development in an *alg-2(0)* background, whereas *air-2* and *chk-1* produced low-penetrance phenotypes in both life histories in an *alg-2(0)* background.

Prior to this work, the only kinase-encoding gene known to enhance *alg-1(0)* phenotypes after dauer was *kin-3,* used as a positive control in this study (Alessi *et al*. 2015). During continuous development, three kinases have been shown to phosphorylate miRISC components to modulate different aspects of miRNA activity: KIN-1/PKA, casein kinase 1 (CK1) and CK2 (Alessi *et al*. 2015; Huberdeau *et al*. 2022; Shah *et al*. 2023). In our screen, only the CK2-encoding genes *kin-3* and *kin-10* were found to be necessary to prevent vulval defects in *alg-1(0)* mutants after dauer, whereas RNAi of *kin-1* and *kin-19,* encoding PKA and CK1 respectively, did not cause vulval defects in our primary screen (*kin-1,* 0/40; *kin-19,* 0/60). While these findings may be explained by incomplete knockdown of *kin-1* and *kin-19,* it is also possible that distinct kinases modulate miRISC in the dauer life history. This study lays the foundation for future studies that can dissect the molecular relationship between these genes and miRISC in *C. elegans* and other species.

## Data availability

All *C. elegans* strains and newly made RNAi clones are available upon request. The authors state that all data necessary for confirming the conclusions of the article are present within the article, figures, and tables, including supplementary information.

## Acknowledgments

We thank Jennifer Schisa, and Steven Gorsich, and Karp lab members (Central Michigan University) for their helpful advice throughout this project. We are grateful to BIO 546 students (Central Michigan University) who helped with pilot experiments. We thank WormBase. Some strains were provided by the CGC, which is funded by NIH Office of Research Infrastructure Programs (P40 OD010440). This project was funded by CAREER 1652283 to XK from the National Science Foundation.

## Notes

### Competing Interest Statement

The authors have declared no competing interest.

### Summary of Updates

Some additional detail was added to the Introduction and Discussion sections and small typos were corrected.

