## Supplemental Figures 1 and 2 for "An RNAi screen for conserved kinases that enhance microRNA activity after dauer in *C. elegans*"

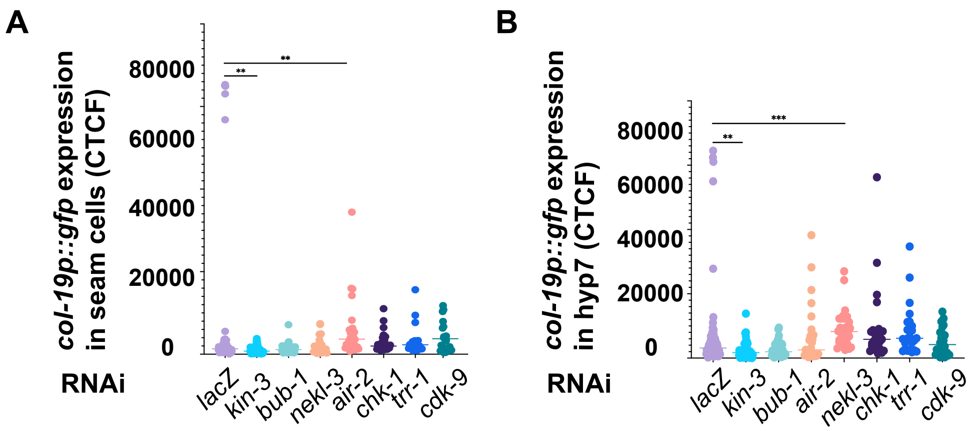


**Figure S1**. **RNAi of six kinase-encoding genes identified in the primary screen did not reduce *col-19p::gfp* after dauer.** The corrected total cell fluorescence (CTCF) value of *col-19p::gfp* expression in seam cells **(A)** or hyp7 nuclei **(B)** in post-dauer adult animals, as quantified using ImageJ. The CTCF value was calculated by subtracting the mean fluorescence background from the integrated density. Only the *kin-3* control showed reduced *col-19p::gfp* expression in either seam cells or hyp7, compared to *lacZ.* RNAi of *air-2* and *nekl-3* produced slightly increased *col-19p::gfp* expression in seam cells or hyp7, respectively. ** $p$-value < 0.004, *** $p < 0.0008$, Kruskal-Wallis and Dunn's multiple comparisons test. Note that when the outliers from the *lacZ* control were removed, *kin-3(RNAi)* treated post-dauer adults still showed significantly reduced *col-19p::gfp* compared to those treated with *lacZ* RNAi.

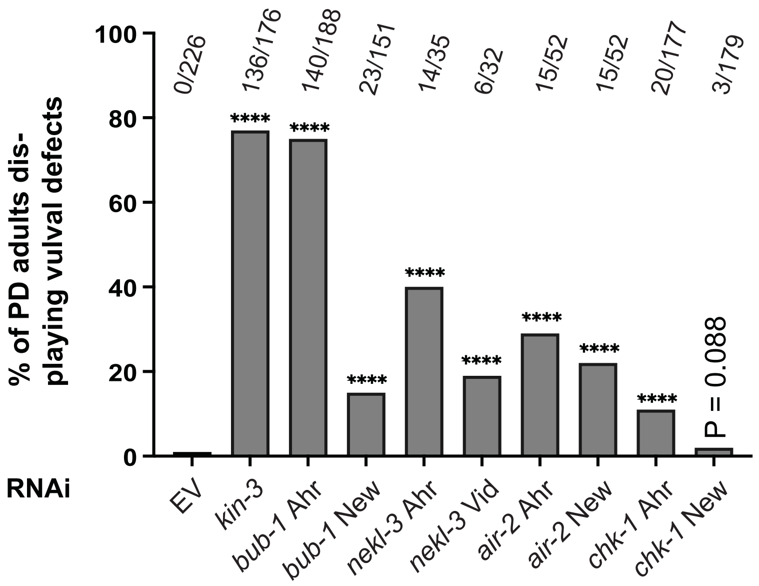


**Figure S2**. **Two independent RNAi clones of kinases identified in the screen produce vulval defects**. For each gene identified as a hit in the primary and secondary screens, a second RNAi clone was tested for its ability to produce vulval defects (Rup and/or Pvl) in young post-dauer *alg-1(0)* adults. Where available, the second RNAi clone was taken from existing RNAi libraries, "Ahr" = Ahringer library, "Vid" = Vidal library (Kamath and Ahringer 2003; Rual *et al.* 2004). Where unavailable, new RNAi clones were created, "New" (see Methods). *p*-value <0.0001 (****), Fisher exact test compared to EV (empty vector) control.
